# Differential nanoscale organization of excitatory synapses onto excitatory vs inhibitory neurons

**DOI:** 10.1101/2023.09.06.556279

**Authors:** Poorna A. Dharmasri, Aaron D. Levy, Thomas A. Blanpied

## Abstract

A key feature of excitatory synapses is the existence of subsynaptic protein nanoclusters whose precise alignment across the cleft in a trans-synaptic nanocolumn influences the strength of synaptic transmission. However, whether nanocolumn properties vary between excitatory synapses functioning in different cellular contexts is unknown. We used a combination of confocal and DNA-PAINT super-resolution microscopy to directly compare the organization of shared scaffold proteins at two important excitatory synapses – those forming onto excitatory principal neurons (Ex→Ex synapses) and those forming onto parvalbumin-expressing interneurons (Ex→PV synapses). As in Ex→Ex synapses, we find that in Ex→PV synapses presynaptic Munc13-1 and postsynaptic PSD-95 both form nanoclusters that demonstrate alignment, underscoring synaptic nanostructure and the trans-synaptic nanocolumn as conserved organizational principles of excitatory synapses. Despite the general conservation of these features, we observed specific differences in the characteristics of pre-and postsynaptic Ex→PV nanostructure. Ex→PV synapses contained larger PSDs with fewer PSD-95 NCs when accounting for size than Ex→Ex synapses. Furthermore, the PSD-95 NCs were larger and denser. The identity of the postsynaptic cell also had a retrograde impact on Munc13-1 organization, as Ex→PV synapses hosted larger Munc13-1 puncta that contained less dense but larger and more numerous Munc13-1 NCs. Moreover, we measured the spatial variability of trans-synaptic alignment in these synapse types, revealing protein alignment in Ex→PV synapses over a distinct range of distances compared to Ex→Ex synapses. We conclude that while general principles of nanostructure and alignment are shared, cell-specific elements of nanodomain organization likely contribute to functional diversity of excitatory synapses. Understanding the rules of synapse nanodomain assembly, which themselves are cell-type specific, will be essential for illuminating brain network dynamics.

## Introduction

Many synaptic proteins are organized into subsynaptic nanoclusters (NCs) to enable their vital functions in information transmission (1). This is true of presynaptic release-related proteins such as Munc13-1, which forms NCs that mark synaptic vesicle release sites within the active zone, and whose density within NCs contributes to variability in probability of release (2–5). Similarly, postsynaptic scaffolds such as PSD-95 form NCs that influence glutamate receptor distribution (6–8), highlighting the importance of subsynaptic protein organization in setting cis-synaptic properties. The impact of nanostructure also extends across the synaptic cleft – for instance, Munc13-1 NCs also contribute to synaptic functions that arise from trans-synaptic relationships, including the magnitude and variance of postsynaptic response amplitudes (9). These trans-synaptic functional impacts likely arise because NCs can be aligned across the synapse to form the trans-synaptic nanocolumn (10), a structural configuration that aligns release sites with glutamate receptors and thus strengthens the postsynaptic response to action potential-evoked neurotransmitter release (7, 10–13. Indeed, dispersal of AMPARs out of the nanocolumn and into the rest of the synapse is sufficient to greatly reduce the amplitude of action potential-evoked postsynaptic currents (14), underscoring the relationship between the nanocolumn and synaptic transmission.

Protein nanoclustering is a synapse architectural feature that has been observed with diverse imaging modalities in myriad preparations, including synaptosomes, dissociated cultures, *ex vivo* slices, and *in vivo* imaging (15–20). However, nanocolumn alignment has been less frequently measured. Nanocolumns of Unc13A and clustered receptors have been identified at the Drosophila neuromuscular junction (21) and alignment of RIM and GABA_A_Rs has been reported in inhibitory synapses (22), suggesting the nanocolumn is an evolutionarily old and widespread structure. Despite this, we lack an understanding of how nanocolumn architecture may vary, even between excitatory synapse types, leaving unresolved whether or which properties of the nanocolumn are conserved. The near ubiquity of core proteins, including Munc13-1 and PSD-95, at different excitatory synapse types suggests their nanostructural relationship might be stereotyped. However, while core proteins are shared, different neuron types express distinct sets of synaptic receptors, adhesion proteins, and accessory proteins (23–25) that could modify the organization of core elements. Thus, cellular context may be a critical variable that alters nanocolumn properties, tuning alignment to support the physiological functions of a particular circuit.

Most explorations of excitatory synapse nanostructure have been conducted on glutamatergic synapses forming onto spiny, excitatory neurons. To test whether cellular context impacts nanostructure, we considered another particularly important excitatory synapse type, the afferent excitatory synapses onto parvalbumin-expressing hippocampal interneurons (PV-INs). Excitatory drive onto PV-INs mediates numerous critical aspects of cognition and pathology (26, 27), so understanding their synaptic molecular architecture is of utmost importance. Further, PV-INs express a profile of proteins that is relatively consistent across brain regions (24) but differs from that of principal cells (28–30), making them an ideal candidate for testing the conservation of nanocolumn properties.

Here we used confocal imaging and 3D DNA-PAINT super-resolution microscopy to investigate differences in protein spatial organization between excitatory synapses onto CaMKII-expressing excitatory neurons and onto PV-INs. This direct, quantitative comparison revealed that the generalities of synaptic nanostructure and the nanocolumn are conserved between these two synapse types. However, postsynaptic cell identity impacted the specific nanoscale organization of both postsynaptic and presynaptic proteins, as well as the trans-synaptic spatial relationship between them. Our results underscore that while being generally conserved, subsynaptic nanostructure may represent an additional axis upon which synaptic transmission can be tuned to fit the needs of specific cells.

## Results

### Pre and postsynaptic protein abundance and density depend on postsynaptic cell type

To measure protein organization at specific synapse types, we identified postsynaptic cells as either excitatory neurons or PV-INs in DIV21 dissociated rat hippocampal cultures by infecting with lentivirus expressing EGFP under a CaMKII promoter or by immunostaining for PV, respectively. The robust expression of either EGFP or PV throughout the soma and dendrites allowed clear delineation of expected cell morphologies: compared to excitatory neurons, PV-INs tended to have larger somata and longer dendrites with a distinct, near-total absence of dendritic spines (31). All cells were also immunostained for presynaptic Munc13-1 and postsynaptic PSD- 95 to identify afferent excitatory synapses forming onto either the excitatory neurons (Ex→Ex) or PV-INs (Ex→PV).

Ex→PV synapses qualitatively appeared larger and brighter than their Ex→Ex counterparts by confocal microscopy and had a ‘railroad track’-like organization, appearing in parallel along the dendrite (Fig 1A-B). To achieve an unbiased quantification of synapse properties, we built a semi-automated workflow centered around the puncta detection tool SynQuant (31; Fig S1). Using this approach, we were able to quantify 40,751 and 26,627 synapses from 48 CaMKII-expressing neurons and 27 PV-INs, respectively, across three independent cultures.

**Figure 1.**
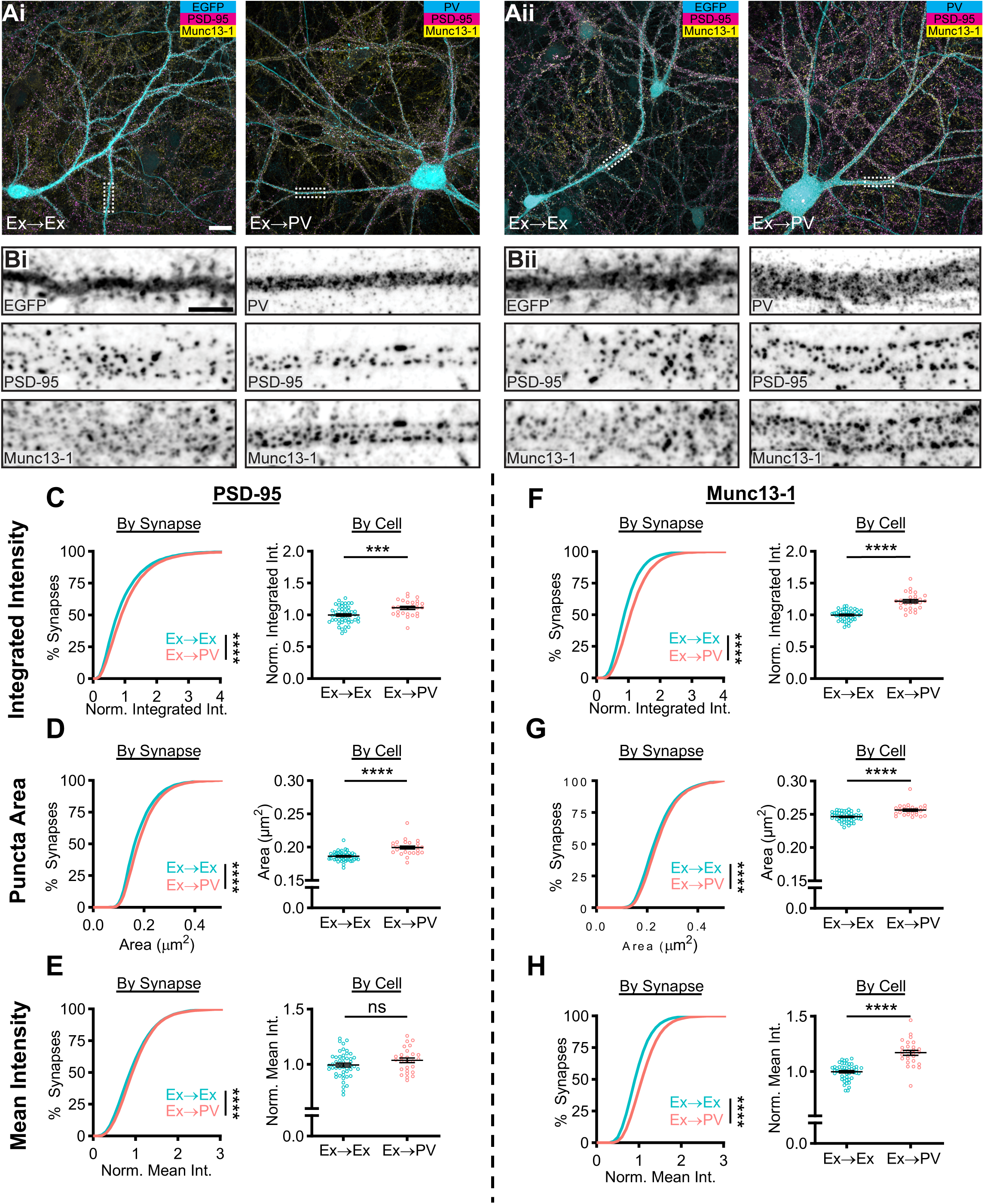
PV-INs host larger excitatory synapses with higher Munc13-1 density than CaMKII-expressing neurons. **A)** Example neurons in dissociated hippocampal cultures from two independent cultures (i, ii). Excitatory synapses formed onto excitatory neurons (Ex→Ex) were identified by CaMKII promoter-driven EGFP expression (Ai-ii, left), while excitatory synapses formed onto PV-INs (Ex→PV) were identified by PV staining (Ai-ii, right). Scale bar: 20 µm. **B)** Representative dendrite stretches from example cells above showing, from top to bottom, cell identifying marker, PSD-95 staining, and Munc13-1 staining. Scale bar: 5 µm. **C-H)** Analyses of synaptic protein content and puncta size. All data are presented on a per synapse (left) and per cell (right) basis. Per synapse analyses, presented in cumulative distributions, represent the direct observations (Ex→Ex, n = 40,751 synapses; Ex→PV, n = 26,627 synapses; p<0.0001 for all). Some CDFs are cut off at the 99^th^ percentile for visibility. Per cell analyses were conducted to account for cell-to-cell variability (Ex→Ex, n = 48 cells; Ex→PV, n = 27 cells; statistics below). Intensity measures are normalized within culture week to the average of the Ex→Ex group, and each cell is presented as a point along with the mean ± SEM, which is also reported below. **C)** Ex→PV synapses contain more PSD-95 (per cell: Ex→Ex, 1.00 ± 0.019; Ex→PV, 1.11 ± 0.023; p=0.0004). **D)** Ex→PV PSDs are larger than their Ex→Ex counterparts (per cell: Ex→Ex, 0.19 ± 0.0010 µm^2^; Ex→PV, 0.20 ± 0.0021 µm^2^; p<0.0001). **E)** PSD-95 density is not different between synapse types when accounting for cell-to-cell variability (per cell: Ex→Ex, 1.00 ± 0.017; Ex→PV, 1.04 ± 0.021; p=0.1281). **F)** Ex→PV synapses contain more Munc13-1 (per cell: Ex→Ex, 1.00 ± 0.012; Ex→PV, 1.22 ± 0.025; p<0.0001). **G)** AZs are larger in Ex→PV synapses (per cell: Ex→Ex, 0.25 ± 0.0010 µm^2^; Ex→PV, 0.26 ± 0.0016 µm^2^; p<0.0001). **H)** Ex→PV synapse AZs have a greater Munc13-1 density (per cell: Ex→Ex, 1.00 ± 0.011; Ex→PV, 1.17 ± 0.022; p<0.0001).

We first quantified the total amount of PSD-95 at each synapse by measuring its integrated intensity, normalizing within each culture replicate to the mean value at Ex→Ex synapses. The cumulative distribution was right-shifted in PV-INs (Fig 1C, left), indicating more PSD-95 per synapse. As synaptic content can differ based on activity history, we tested if cellular context influenced this shift. Ex→PV synapses still had a higher PSD-95 integrated intensity when averaged per cell (Fig 1C, right). The difference in integrated intensity could be due to either a change in synapse size or protein density, so we next quantified the area and mean intensity of PSD-95 puncta. We found that PSD-95 puncta were larger in Ex→PV synapses when analyzed per-synapse or per-cell (Fig 1D). In contrast, while the mean intensity of PSD-95 in all Ex→PV synapses showed a small increase, the per cell analysis indicated this was not a significant difference (Fig 1E). These results indicate that PV-INs have larger, but not denser, excitatory postsynapses than CaMKII-expressing excitatory neurons.

We next asked whether postsynaptic cell identity was associated with differences in the corresponding afferent active zones (AZ), identified with Munc13-1 immunostaining. The normalized integrated intensity and area of Munc13-1 puncta were each significantly larger at Ex→PV synapses (Fig 1F-G). Further, unlike for PSD-95, the normalized mean intensity of Munc13-1 was higher onto PV-INs (Fig 1H), indicating a greater Munc13-1 density within these AZs. These data together demonstrate that postsynaptic cell identity impacts both pre-and postsynaptic content of core excitatory synapse proteins. The differences in synapse size and Munc13-1 density between these synapse types suggest the potential for differential underlying nanoscale organization of core synaptic proteins.

### PSD-95 nanostructure is differentially organized depending on postsynaptic cell identity

We next used 3D DNA-PAINT to measure whether protein nano-organization differs between Ex→Ex and Ex→PV synapses (Fig 2A-B), focusing first on the PSD. Our super-resolution data showed Ex→PV PSDs were nearly two times the volume of Ex→Ex PSDs (Fig 2C), consistent with the larger puncta area detected with confocal imaging. As PSD-95 NC number scales with synapse size at Ex→Ex synapses (15), we predicted that Ex→PV PSDs would contain more PSD- 95 NCs. To test this, we detected PSD-95 NCs using DBSCAN. We were surprised to see no difference in the average number of PSD-95 NCs per synapse (Fig 2D). This could be due to a different relationship between PSD size and NC number at Ex→PV synapses. While the rate at which NC number increases with PSD size is not different between the two synapse types, Ex→PV synapses notably have fewer PSD-95 NCs for a synapse of a given size (Fig 2E). These results suggest Ex→PV PSDs are not simply enlarged versions of Ex→Ex PSDs but differ in their nano-organizational principles.

**Figure 2.**
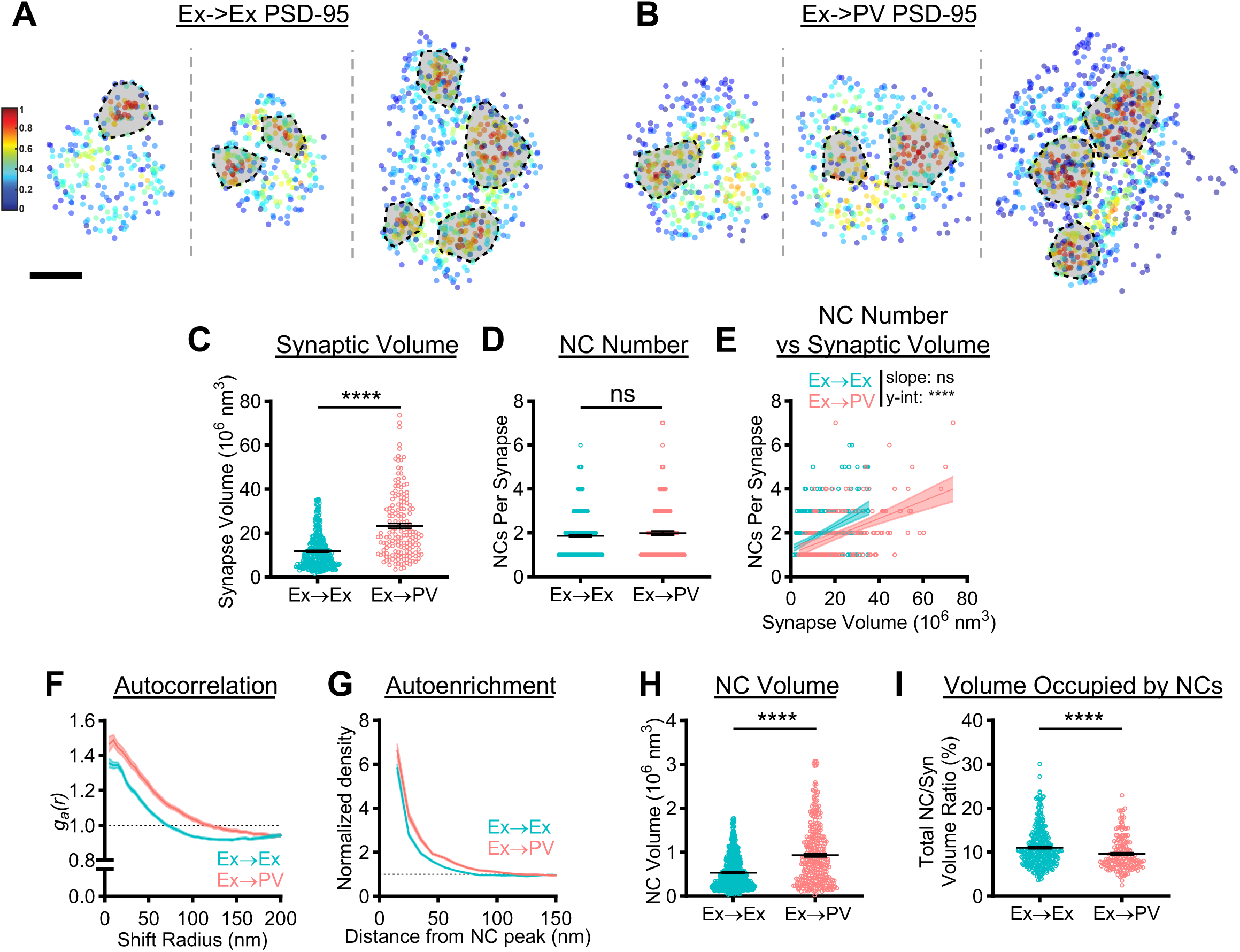
PSD-95 nanoscale organization depends on postsynaptic cell identity. **A, B)** Examples of 3D DNA-PAINT PSD-95 synaptic clusters from A) Ex→Ex and B) Ex→PV synapses. Heat maps indicate the peak density-normalized local density. Black-bordered shapes indicate detected NCs. Scale bar: 100 nm. Parentheticals below indicate mean ± SEM. **C)** PSDs are larger at Ex→PV synapses (Ex→Ex, 11.76 ± 0.41 x 10^6^ nm^3^; Ex→PV, 23.32 ± 1.14 x 10^6^ nm^3^; p<0.0001). **D)** Both synapse types contain the same number of PSD-95 NCs on average (Ex→Ex, 1.87 ± 0.054 NCs/synapse; Ex→PV, 1.99 ± 0.092 NCs/synapse; p=0.4606). **E)** Ex→PV PSDs contain fewer NCs than expected when accounting for PSD size (Ex→Ex, slope: 0.053, y-intercept: 1.25, R^2^: 0.17; Ex→PV, slope: 0.040, y-intercept: 1.058, R^2^: 0.24. slope p=0.1009, y-intercept p<0.0001). **F)** Autocorrelation predicts PSD-95 forms larger, denser NCs in PV cells. **G)** The average Ex→PV PSD-95 NC has a denser and broader PSD-95 distribution. **H)** PSD-95 NC volume is larger at Ex→PV synapses (Ex→Ex, 0.54 ± 0.014 x 10^6^ nm^3^; Ex→PV, 0.94 ± 0.038 x 10^6^ nm^3^; p<0.0001). **I)** Less synapse volume is occupied by PSD-95 NCs at Ex→PV synapses (Ex→Ex, 11.03 ± 0.23% volume; Ex→PV, 9.62 ± 0.29% volume; p<0.0001). Data in C-E and I are individual synapses, and in H individual NCs, with lines at mean ± SEM. Data in E are plotted with line of best fit ± 95% confidence interval. Data in F and G show a line connecting the mean of each bin ± SEM shading. For Ex→Ex, n = 372 synapses or 667 NCs; for Ex→PV, n = 168 synapses or 327 NCs throughout.

To directly interrogate whether PSD-95 nanocluster properties differ between the two synapse types, we first quantified the spatial heterogeneity of PSD-95 within single synapses using a normalized autocorrelation analysis, as described previously (10, 14). The autocorrelation of PSD-95 at Ex→PV synapses was higher in magnitude over longer shift radii, predicting larger, denser PSD-95 NCs than in Ex→Ex synapses (Fig 2F). An autoenrichment analysis, which measures the normalized density of a protein surrounding its NC peak, showed both a higher peak density and broader distribution (Fig 2G) and PSD-95 NC volume was ∼75% larger at Ex→PV synapses (Fig 2H), confirming the autocorrelation predictions. We wondered if the larger PSD-95 NCs at Ex→PV synapses ensured the same fraction of the PSD was occupied by PSD- 95 NCs despite being fewer in number. We found instead that PSD-95 NCs occupied a smaller proportion of the PSD at Ex→PV synapses than at Ex→Ex synapses (Fig 2I). Together, these data demonstrate that PSD-95, a core scaffold protein of excitatory synapses, is differentially organized depending on postsynaptic cell identity. The unique subsynaptic pattern of PSD-95 may strengthen the retention of mobile glutamate receptors within these larger synapses (see Discussion).

### Postsynaptic cell type has retrograde effects on active zone nanostructure

Given the higher Munc13-1 density at Ex→PV synapses (Fig 1J), we asked whether the postsynaptic cell identity impacted the afferent Munc13-1 nanoscale organization (Fig 3A-B). Consistent with our confocal imaging observations, Munc13-1 occupied a larger total volume at Ex→PV than at Ex→Ex synapses (Fig 3C), suggestive of a larger active zone. Unlike PSD-95 NCs, Munc13-1 NCs were more numerous in Ex→PV synapses (Fig 3D) suggesting these AZs contain more release sites (an average of ∼1 more per synapse). In hippocampal CA1 pyramidal neuron synapses onto fast-spiking interneurons (FSINs), the number of Munc13-1 nanoclusters scales with active zone size yet is strikingly variable even in active zones of similar size (3). We tested whether this was true both in Ex→PV and Ex→Ex synapses. The number of Munc13-1 NCs were similarly correlated with the size of the AZ in each synapse type (Fig 3E). The spread of the data in the regression analysis is indicative of large variability in Munc13-1 NC number across similarly sized AZs. Agnostic to synapse size, the variability in NC number per synapse was large but not different between groups (Ex→Ex: CV 0.58 vs Ex→PV: CV 0.59). Together, these data are consistent with previous findings and suggest that postsynaptic cell identity can influence AZ size.

**Figure 3.**
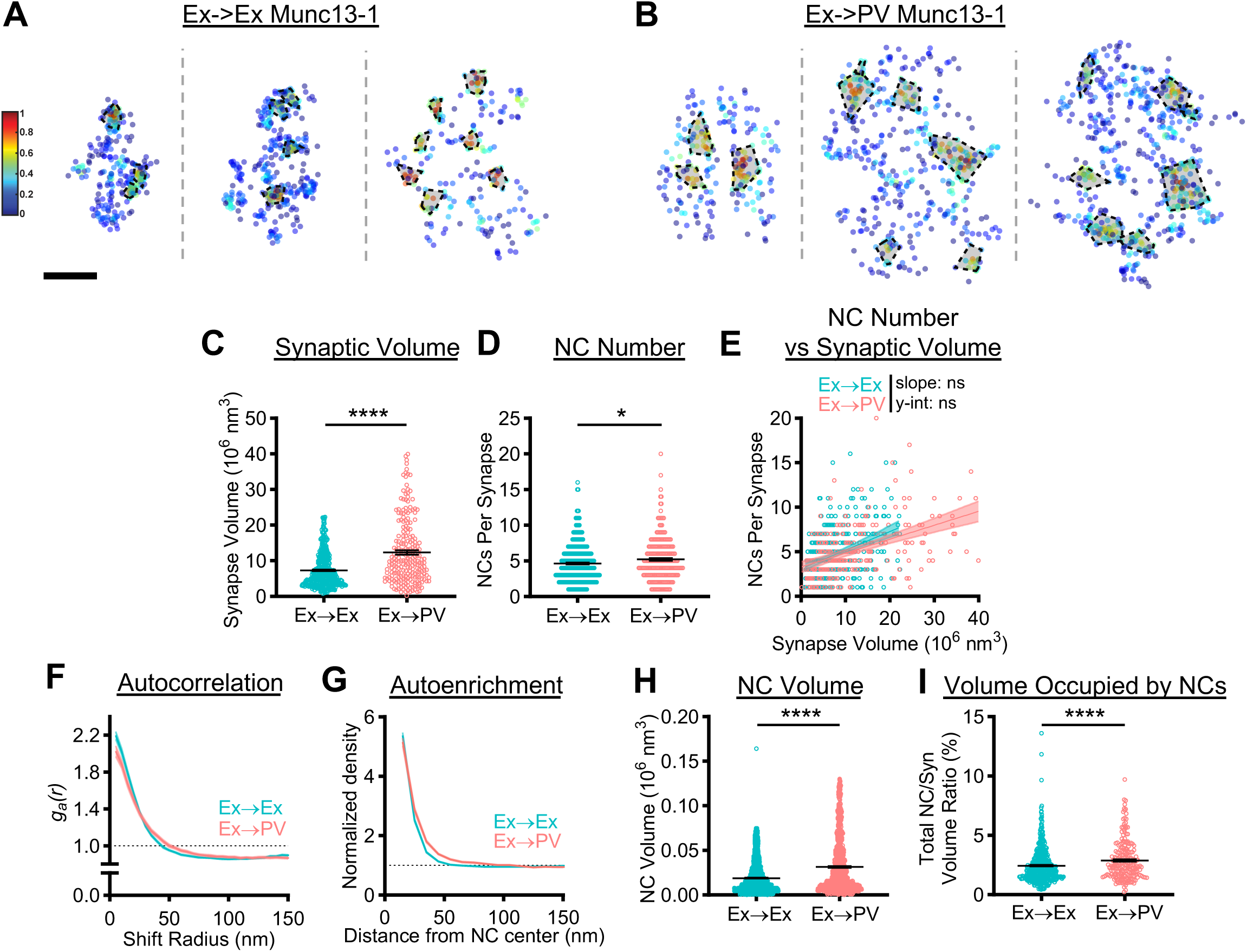
Postsynaptic cell identity has a trans-synaptic influence on active zone Munc13-1 nanostructure. **A, B)** Examples of 3D DNA-PAINT Munc13-1 synaptic clusters from A) Ex→Ex and B) Ex→PV synapses. Heat maps indicate peak density-normalized local density. Black-bordered shapes indicate detected NCs. Scale bar: 100 nm. Parentheticals below indicate mean ± SEM. **C)** Munc13-1 AZ clusters are larger in Ex→PV synapses (Ex→Ex, 7.31 ± 0.22 x 10^6^ nm^3^; Ex→PV, 12.34 ± 0.59 x 10^6^ nm^3^; p<0.0001). **D)** Ex→PV synapses contain more Munc13-1 NCs. (Ex→Ex, 4.63 ± 0.13 NCs/synapse; Ex→PV, 5.21 ± 0.20 NCs/synapse; p=0.0143). **E)** Munc13-1 NC number scales similarly with AZ size between synapse types (Ex→Ex, slope: 0.21, y-intercept: 3.10, R^2^: 0.14; Ex→PV, slope: 0.16, y-intercept: 3.28, R^2^: 0.21; slope p=0.1016, y-intercept p=0.1887). **F)** Autocorrelation predicts Munc13-1 forms larger, less dense NCs at Ex→PV synapses. **G)** The average Ex→PV Munc13-1 NC has a broader Munc13-1 distribution with lower central density. **H)** Munc13-1 NC volume is larger at Ex→PV synapses (Ex→Ex, 0.019 ± 0.00042 x 10^6^ nm^3^; Ex→PV, 0.032 ± 0.00096 x 10^6^ nm^3^; p<0.0001). **I)** More of the synapse volume is occupied by Munc13-1 NCs in Ex→PV synapses (Ex→Ex, 2.45 ± 0.071% volume; Ex→PV, 2.89 ± 0.11 % volume; p<0.0001). Data in C-E and I are individual synapses, and in H individual NCs, with lines at mean ± SEM. Data in E plotted with line of best fit ± 95% confidence interval. Data in F and G show a line connecting the mean of each bin ± SEM shading. For Ex→Ex, n = 454 synapses or 1839 NCs; for Ex→PV, n = 229 synapses or 1046 NCs throughout.

A closer look at Munc13-1 nanocluster properties revealed further postsynaptic cell type-dependent differences. Autocorrelation analysis of the Munc13-1 distribution in Ex→PV synapses showed a lower initial magnitude with a concomitant rightward shift in the middle portion of the curve (Fig 3F), predicting larger, but less dense NCs than at Ex→Ex synapses. Consistent with these predictions, Munc13-1 NCs at Ex→PV synapses had higher local density at longer distances from the NC peak (Fig 3G) and were larger (Fig 3H) than those at Ex→Ex synapses. The combination of more numerous and larger Munc13-1 NCs at Ex→PV synapses resulted in a larger fraction of the AZ being occupied by these NCs (Fig 3I). These data together demonstrate that the postsynaptic cell influences apposed AZ size as well as Munc13-1 NC volume and density, parameters which may affect synaptic vesicle docking, priming, or probability of release, which are high at Ex→PV synapses (33–36).

### Impact of postsynaptic cell type on trans-synaptic alignment

In the prototypical Ex→Ex synapse, synaptic strength depends on the trans-synaptic coordination of nanostructure (11, 14, 37). Given that in Ex→PV synapses more of the AZ is occupied by Munc13-1 NCs but less of the PSD is occupied by PSD-95 NCs (compare Figs 2I and 3I), we predicted that trans-synaptic alignment in Ex→PV synapses would be weaker. We first measured the distance between nearest NC peaks of opposing proteins (Fig 4A and 4B). Consistent with larger synapses, in Ex→PV synapses the peak of Munc13-1 NCs were slightly further away from the nearest identified PSD-95 NC peak, and vice versa (Fig 4C), suggesting weaker alignment between release sites and postsynaptic scaffolds. However, while the NC peaks may be further apart, NCs, particularly of PSD-95, do not contain the majority of the scaffold protein (7), limiting the utility of the peak-to-peak analysis. We thus analyzed alignment with two additional approaches that consider protein distribution without relying on identifying NCs of each.

**Figure 4.**
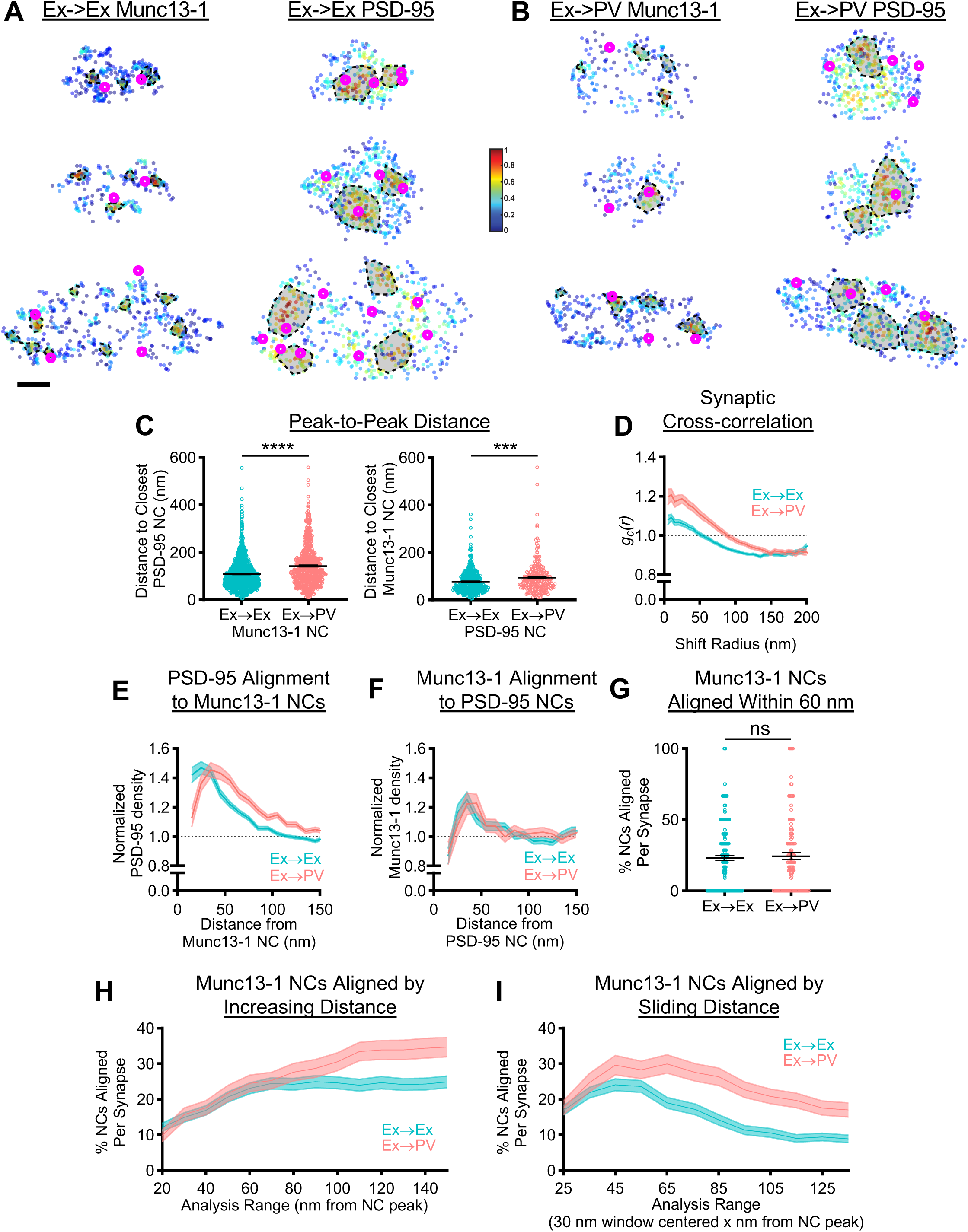
Postsynaptic cell identity impacts the characteristic trans-synaptic nanoscale relationship between Munc13-1 and PSD-95. **A, B)** Example 3D DNA-PAINT synapses from A) Ex→Ex and B) Ex→PV synapses. Magenta circles indicate centers of NCs of opposing protein and heat map indicates peak density-normalized local density of labelled protein. Each row is an individual synapse with corresponding AZ and PSD clusters in the indicated columns. Scale bar: 100 nm. Parentheticals below indicate mean ± SEM. **C)** Munc13-1 and PSD-95 NC peaks are further from each other in Ex→PV synapses (Munc13-1 NC to closest PSD-95 NCs: Ex→Ex, 108.7± 1.90 nm, n = 1141 NCs; Ex→PV, 142.9 ± 3.21 nm, n = 695 NCs; p<0.0001. PSD-95 NC to closest Munc13-1 NCs: Ex→Ex, 77.09 ± 2.21 nm, n = 448 NCs; Ex→PV, 93.82 ± 4.28 nm, n = 246 NCs; p=0.0004). **D)** Synaptic cross-correlation indicates stronger trans-synaptic alignment in Ex→PV synapses (Ex→Ex, n = 245 synapses; Ex→PV, n = 129 synapses). **E)** The enrichment of PSD-95 density across from Munc13-1 NCs indicates an offset in nanocolumn alignment in Ex→PV synapses (Ex→Ex, n = 1067 NCs; Ex→PV, n = 623 NCs). **F)** The enrichment of Munc13-1 density across from PSD-95 NCs has a broader, rightward-skewed peak in Ex→PV synapses (Ex→Ex, n = 474 NCs; Ex→PV, n = 263 NCs). **G)** Both synapse types have equal proportions of Munc13-1 NCs aligned to PSD- 95 within 60 nm of the NC peak (Ex→Ex, 23.11 ± 1.57%; Ex→PV 24.49 ± 2.35%, p=0.7837). **H)** Both synapse types have an equal proportion of Munc13-1 NCs aligned to PSD-95 up to 100 nm away from the NC peak, after which they diverge. **I)** Ex→PV synapse Munc13-1 NCs are aligned at a higher percentage over a broader range than their Ex→Ex counterparts (for G-I, Ex→Ex, n = 237 synapses; Ex→PV, n = 124 synapses). Data in C are individual nanoclusters with lines at mean ± SEM. All other data are synapses. Data in D-F and H-I show a line connecting mean of each bin ± SEM shading.

To dispense with NC identification entirely, we first used a cross-correlation analysis that reports the overall similarity of nanoscale protein distributions within the synapse (10). This showed a higher magnitude over shift radii up to 150 nm at Ex→PV synapses (Fig 4D), consistent with stronger trans-synaptic nanoscale alignment of Munc13-1 and PSD-95 overall. The increase in the width of the cross-correlation is consistent with the larger NCs found in Ex→PV synapses, since in a synapse displaying nanocolumn alignment this would position more protein near opposing areas of high protein density, a fact not captured in the peak-to-peak distances.

To test alignment across from identified NCs of just one protein species, we measured the cross-enrichment, which is the normalized radial distribution of the opposing protein across from a given NC’s peak. PSD-95 was strongly cross-enriched to Munc13-1 NC peaks, and vice versa, within both synapse types (Fig 4E-F), suggesting that despite larger and slightly broader spaced NCs, the overall relationship of trans-synaptic spatial alignment is maintained in Ex→PV synapses. Along with the cross-correlation analysis, this confirms the presence of a conserved nanocolumn architecture between Ex→Ex and Ex→PV synapses.

Intriguingly, however, the cross-enrichment analyses revealed specific organizational differences between the two synapse types. The peak density of PSD-95 was shifted 10 to 20 nm further away from Munc13-1 NCs in Ex→PV synapses (Fig 4E). The offset was also reflected in the cross-enrichment of Munc13-1 density to PSD-95 NCs, which was overall extremely similar between the two synapse types but showed a broader, rightward-skewed peak in Ex→PV synapses (Fig 4F). These data suggest that nanocolumns within Ex→PV synapses comprise a slightly offset or perhaps annular trans-synaptic organization when compared to Ex→Ex synapses. This is an important notion because the most widely considered working model of nanocolumn alignment is that cell adhesion molecules (CAMs) with discrete interactions across the cleft provide the basis for a stereotyped transcellular architecture. Since different cells and synapses express distinct subsets of CAMs that may interact with distinct molecular populations, this raises the possibility that the nanocolumn in different synapse types could be constructed with a consistent but unique structure. Thus, while nanocolumns exist in both synapses, Ex→PV synapses may contain different nanocolumn ‘subtypes’ than Ex→Ex synapses.

To establish whether the unexpected shift in cross-enrichment in Ex→PV synapses indicated a single population of consistently structured nanocolumns or rather a broad distribution of nanocolumn ’subtypes’, we developed analyses to examine the heterogeneity in alignment in the population of Munc13-1 NCs within the analyzed synapses. We determined the percentage of Munc13-1 NCs that were aligned per synapse by testing whether each Munc13-1 NC was statistically enriched with PSD-95 or not within a specific radius. When we applied this measure using a radius of 60 nm as in previous work (10) and calculated the percent of aligned Munc13-1 NC per synapse, the same percentage were aligned in each synapse type (Fig 4G). However, NCs are larger at Ex→PV synapses, and so to evaluate this relationship more systematically, we varied the width of the analysis range from 20 to 150 nm, calculating for each range the percent of Munc13-1 NCs per synapse statistically aligned to PSD-95. The percentage of aligned Munc13- 1 NCs was similar between Ex→Ex and Ex→PV synapses up to 100 nm away from the NC peak (Fig 4H). Given that the difference in the peak cross-enrichment to Munc13-1 NCs was observed within this distance (Fig 4E), we probed for a specific distance at which Munc13-1 was enriched with PSD-95 using a sliding window of 30 nm for the analysis. This revealed different preferred distances for the two synapse types even within those first 100 nm; most Munc13-1 NC alignment occurred around 35-55 nm in Ex→Ex synapses, but in Ex→PV synapses maximal alignment occurred further away and across a wider range of distances (45-75 nm; Fig 4I). Overall, these results are consistent with the idea that while nanocolumn architecture is generally conserved, Ex→PV synapses host a wider distribution of nanocolumn ‘subtypes’ than Ex→Ex synapses, presumably arising from engagement of their unique set of cleft-resident molecules.

## Discussion

In this study, we used complementary intermediate-scale confocal imaging and single-molecule imaging via DNA-PAINT to directly compare the nanostructural organization of excitatory synapses forming onto PV-INs vs onto excitatory neurons. Our detailed nanostructural analysis indicates that both PSD-95 and Munc13-1 at excitatory synapses on PV-INs form nanoclusters that are aligned across the synapse from one another, suggesting the nanocolumn is a conserved organizational principle of excitatory synapses. This interpretation is suggested by repeated observations of synaptic protein nanoclusters (15–20) and alignment (21, 22) in other synapses and systems, but our detailed comparison with identical analyses at high imaging resolutions provides direct support for this conclusion. Conservation of nanocolumn alignment emphasizes the idea that common structural mechanisms govern the establishment of synaptic strength and provide distinct means for plastic changes at single synapses (1, 38). Indeed, experimental disruption of synaptic cleft interactions perturbs alignment of receptors with release sites and rapidly reduces the strength of synaptic transmission (14, 37, 39), suggesting CAMs play key roles in these mechanisms. Interestingly, while PV-INs express a unique set of CAMs compared to excitatory neurons (28–30) and form synapses on the quite morphologically distinct dendritic shaft instead of the spine head, Ex→PV synapses still form nanocolumns, underscoring the robust nature of this architecture and suggesting it may be widely conserved *in vivo*.

Despite the general conservation of nanostructural features, however, key characteristics of both PSD-95 and Munc13-1 distributions depended on the identity of the postsynaptic cell. PSDs in Ex→PV synapses were larger overall with the same number of NCs, as if they were “stretched” Ex→Ex PSDs. However, individual NCs were larger and denser at Ex→PV synapses. Formation of these larger, denser PSD-95 NCs in PV-INs could be driven by PSD-95 interactors not found in excitatory neurons, such as Btbd11 (40), which might alter scaffold organization. The unique pattern of PSD-95 is expected to alter the distribution and mobility of NMDARs and AMPARs within these synapses as they diffuse within and between scaffold nanoclusters (41, 42). Conversely, PV-INs express a unique complement of Ca^2+^-permeable AMPARs and TARPs (31, 43, 44) that may also exert an instructive influence on scaffold organization. These differences in organizational properties between synapse types suggest PSD-95 is susceptible to organizational influences of other proteins in the synapse. This is in some contrast to the idea that PSD-95 could be a dominant driver of PSD organization (45), potentially as a central player in the phase separation properties of many synaptic proteins examined *in vitro* (46). In a general sense, it seems likely that the unique combination of molecular players in particular synapses could produce distinct classes of subsynaptic protein organization, either through a cooperative phase separation mechanism or through direct binding interactions.

One clear outcome of our work is that the organization of Munc13-1 within the active zone is dependent on postsynaptic cell identity. This key protein regulates both the docking and functional priming of SVs at the presynaptic membrane (47–51), so its distribution within the AZ has important consequences for neurotransmitter release. Presuming the axons in the synapses we analyzed drew equally from any excitatory cell type present in these cultures, these data suggest retrograde regulation of the number and properties of presynaptic Munc13-1 NCs by the postsynaptic cell. It may be that distinct CAM expression in the postsynaptic targets of a given axon (28) instructs afferent organization by selecting for specific presynaptic interacting partners that differentially organize the AZ. Functional impacts of such a mechanism have been described, as release probability from CA1 neurons onto different interneuron types is influenced by postsynaptic expression of neuroligin-3 (52) or ELFN1 (36). Beyond synapse strength, Ex→PV synapses are more stable (53), and we speculate that augmented or specific CAM content could lead both to enhanced alignment and decreased synaptic turnover.

Munc13-1 was present at greater density at Ex→PV synapses by confocal, and DNA-PAINT revealed it was organized into more numerous and larger, but less dense, NCs (Fig 3). The number of Munc13-1 NCs that we observe within an active zone correlates with the number of release sites (5), suggesting these synapses accommodate more docked vesicles at steady state. Presuming the release probability per site within an AZ is equal (3, 5), this difference in nanostructure producing a larger physiological N is expected to lead to a stronger excitatory drive onto PV-INs. The size and density of Munc13-1 nanoclusters identified by immunogold EM varies considerably between AZs of CA1 pyramidal cell→FSIN synapses (3). Here, we extend this observation in finding that the organization of individual Munc13-1 NCs varies between excitatory synapses onto different cell types. However, whether changing properties within single Munc13- 1 NCs affects functions in docking and priming remains an open question of intense interest. Though reducing Munc13 content by knockout alters both processes (47, 50), Karlocai et al. (2021) argue that the high docked vesicle occupancy at release sites (5, 33) suggests that varying Munc13-1 content per release site may principally control release probability at these sites, consistent with a potential effect on vesicle priming (33). Our findings are also consistent with previous work that suggests Schaffer collateral synapses onto CA1 interneurons have a high probability of release due, in part, to a larger readily releasable pool (35). Thus, it is tempting to speculate that the density of Munc13-1 within NCs may be linked with vesicle priming state, but what factors drive the change in Munc13-1 distribution, and the resulting impact on active zone and Ex→PV synapse function, remain to be determined.

The trans-synaptic spatial relationship between Munc13-1 and PSD-95, while conserved, was quantitatively different between Ex→Ex and Ex→PV synapses. In general terms, the alignment was stronger in Ex→PV synapses as revealed by the cross-correlation. This is of interest, since other things being equal, in large synapses some glutamate receptors will lie farther away from some release sites than in small synapses, thus experiencing a lower amplitude glutamate transient. The augmented correlation of release sites and receptor scaffolds we observe would counteract this and suggests that controlling protein organization within the synapse interior to regulate trans-synaptic alignment may reflect an equalizing mechanism in synapse types of varying geometry.

An unexpected and intriguing difference between synapses was the cross-enrichment offset of PSD-95 from Munc13-1 NCs. Further exploration of this offset with a detailed spatial analysis of alignment suggested that Ex→PV synapses contained a wider distribution of nanocolumn ‘subtypes’ than their Ex→Ex counterparts. One interpretation is that the larger PSDs with larger PSD-95 NCs in Ex→PV synapses could represent an expanded platform upon which CAMs can bind, thus enabling more variance in their positioning and, consequently, more variance in the positioning of presynaptic structures around regions of high local PSD-95 density. In this model, Ex→PV synapse Munc13-1 NCs would be assembled at generally more distributed points around PSD-95 NCs. This is supported by the peak-to-peak measurements, the shift in the Munc13-1 NC cross-enrichment, and the breadth of the peak in both the PSD-95 NC cross-enrichment and the Munc13-1 NC sliding window analysis (Fig 4). Manipulating PSD-95 clustering directly via artificial oligomerization to determine the impact on nanocolumn heterogeneity may permit a direct test of this model.

The existence of nanocolumns that are more offset in Ex→PV synapses would predict a displacement of AMPARs away from sites of release, but the average lateral offset (∼10-20 nm) is itself modest and not likely to be functionally impactful. (Note that we did not compare AMPAR distribution directly in these experiments since the subunit composition is not equal and it is currently not possible to determine subtypes in imaging experiments.) Nevertheless, the organization suggests several possible differences in the synaptic mechanisms which control receptor mobility and determine their positioning. The more abundant space between PSD-95 NCs in the relatively enlarged Ex→PV synapses (Fig 2I) suggests that an AMPAR diffusing within a PV-IN PSD experiences more space without an aligned release site. Thus, mechanisms to keep AMPARs in a PSD-95 NC, once one is encountered, could be particularly critical for maintaining the efficacy of the Ex→PV synapse. Indeed, larger, denser PSD-95 NCs may “capture” diffusing AMPARs more readily through their TARP intermediates, especially since PV-INs predominately express stargazin/TARPγ2, which has a higher affinity for PSD-95 than the prevalent TARP in Ex→Ex synapses, TARPγ8 (44). Cleft interactions may play a role as well. Neuronal pentraxins expressed predominantly in PV-INs interact directly with the N-terminal domains of AMPARs (54) and bind to presynaptic neuronal pentraxin receptor (NPR) to help concentrate AMPARs (55, 56). An offset in Munc13-1 and PSD-95 alignment could thus physically reflect or accommodate the presence of NPR in the AZ within the nanocolumn, providing an additional interaction to increase AMPAR dwell time there. Analysis of AMPAR mobility in these cell types following manipulation of either neuronal pentraxins or TARP content may help resolve these mechanisms. In a general sense, the goal of a synapse is to accumulate and stabilize receptors near neurotransmitter release sites, and the density pattern of PSD-95 will determine how receptors are handled in these two synapse types. This might be particularly impactful in PV-INs, where the dendritic shaft may not impose the same constraints to synapse size and receptor mobility as the membrane structure and bending of a dendritic spine.

Our most general conclusion is that rules of nanodomain assembly – arising from cell-type specific expression patterns – will elaborate synapse functional diversity. Expanding the analyses here to other Ex→IN synapse types will be instructive in efforts to understand this diversity. Note that presynaptic protein organization can be seen independently to influence synaptic strength by establishing the number of release sites, release probability, and features of facilitation and depression of release rate during sustained activity, whereas postsynaptic organization affects strength via controlling the mean number and type of receptors to be activated. Critically, trans-synaptic nanoalignment integrates both these influences (9), helping to determine the resultant quantal amplitude for basal activity frequencies but also creating a higher order impact on frequency-dependent information transfer properties. Thus, defining the nanostructural features and relationships among active zone, cleft, and PSD proteins will be essential for relating expression profiles and ultrastructure to functional diversity across the brain.

## Materials and methods

### DNA constructs

pFCaGW is a lentiviral vector that expresses EGFP (G) under the CaMKII promoter (Ca). The pCaMKII-EGFP transcriptional unit was assembled into a custom backbone (derived from (57), a gift from Alexandros Poulopoulos, and the pEGFP-N1 backbone (Clontech)) by Golden Gate Assembly (NEB), with the CaMKII promoter from mCh-GluA1-CIB (a gift from Matthew Kennedy (Addgene plasmid # 89444; http://n2t.net/addgene:89444; RRID:Addgene_89444)), and EGFP from pEGFP-N1. pCaMKII-EGFP was inserted by NEB HIFI Assembly into the PacI site of a modified pFUGW vector (pFW), where the ubiquitin promoter and EGFP were deleted by KLD mutagenesis (NEB), yielding pFCaGW. For all cloning reactions, fragments were generated by PCR with KAPA HiFi HotStart DNA polymerase (Roche) or by DNA synthesis (IDT). psPAX2 (Addgene plasmid # 12260; http://n2t.net/addgene:12260; RRID:Addgene_12260) and pMD2.G (Addgene plasmid # 12259; http://n2t.net/addgene:12259; RRID:Addgene_12259) were gifts from Didier Trono.

### Lentivirus production

HEK293T cells (ATCC CRL-3216) were maintained in DMEM + 10% FBS and penicillin/streptomycin. To make lentivirus, 5 x 10^6^ cells were plated on a 10 cm dish, then transfected 24h later with 6 µg pFCaGW, 4 ug psPAX2, and 2 µg pMD2.G using PEI (1 µg DNA : 3 µg PEI) for 4-6h before replacing the HEK media with neuron culture media. The media was harvested 48h later, centrifuged at 1000 x g for 5 minutes and filtered through a 0.45 µm PES filter to remove debris, and stored at -80˚C in single-use aliquots. Titers were routinely in the 10^5^- 10^6^ IFU/mL range.

### Primary neuron culture

All animal procedures were approved by the University of Maryland Animal Use and Care committee. Dissociated primary hippocampal neuron cultures were prepared from E18 Sprague-Dawley rat embryos (Charles River) of both sexes. Hippocampus was isolated and dissociated with trypsin, and cells were plated on poly-L-lysine-coated coverslips at 30,000 cells/coverslip (18 mm #1.5, Warner Instruments) in Neurobasal A + GlutaMax, gentamycin, B27 supplement, and 5% FBS. After 24 hours, the media was changed to the same but lacking FBS, and after 1 week supplemented with an additional half volume media + FUDR to suppress glial growth. Neurons were infected at DIV5 with 50 µl of unconcentrated pFCaGW lentivirus and fixed at DIV21 for both confocal and DNA-PAINT experiments.

### Antibody-dye conjugation

300 µl of donkey anti-mouse IgG2a was concentrated on a 100K MWCO Amicon spin filter (Sigma) and diluted to ∼2.5 mg/mL with pH8.3 PBS, then mixed with NHS-Cy3B (GE) at ∼13:1 molar ratio of dye:IgG for 1h at RT to achieve a final dye/IgG ratio of ∼3:1. Excess dye was removed by Zeba 7K MWCO spin desalting column (Thermo Fisher), and the dye/protein ratio confirmed by absorbance at 559 nm and 280 nm according to the manufacturer’s instructions. Conjugated antibody was diluted to ∼1.25 mg/mL in 50% glycerol, aliquoted, and stored at -20˚C.

### Immunostaining

pFCaGW-infected and naïve coverslips from the same plate were fixed with 2% PFA + 4% sucrose in PBS for 10 minutes at room temperature (RT), washed 3 x 5 minutes with PBS + 100 mM glycine (PBSG), permeabilized 20 minutes RT with 0.3% Triton X-100 in PBSG, and blocked 1h RT with 10% donkey serum + 0.2% Triton X-100 in PBSG.

For confocal imaging, the neurons were stained overnight at 4˚C with primary antibodies mouse IgG2A anti-PSD-95, rabbit anti-Munc13-1, and chicken anti-GFP (for pFCaGW-infected cells) or guinea pig anti-PV (for naïve cells) in 5% donkey serum + 0.1% Triton X-100 in PBSG. The next day, cells were washed 3 x 5 minutes in PBSG, then incubated with the secondary antibodies goat anti-mouse IgG2a Cy3B, donkey anti-rabbit AlexaFluor647, and either donkey anti-chicken AlexaFluor488 (pFCaGW) or donkey anti-guinea pig AlexaFluor488 (naïve) for 1 hour at RT in PBSG. Finally, the cells were washed 3 x 5 minutes PBSG, post-fixed with 4% PFA + 4% sucrose in PBS for 15 minutes, and washed 3 x 5 minutes PBSG again before storage at 4˚C until imaging.

For DNA-PAINT, PSD-95 and Munc13-1 were stained with primaries preincubated with custom-made single-domain antibodies (sdAbs; Massive Photonics) carrying one of two oligonucleotide docking strands optimized for DNA-PAINT, as described (58, 59). Briefly, the primary antibodies against PSD-95 and Munc13-1 were incubated separately with a 2.5-fold molar excess of anti-mouse sdAb-F1 or anti-rabbit sdAb-F3, respectively, for 20 minutes at RT, to saturate the antibody with sdAb. Rabbit Fc fragment was added to the Munc13-1 incubation at 2-fold molar excess for a further 20 minutes to remove unbound nanobody. Both preincubations were then diluted to their final working concentrations in 5% donkey serum + 0.1% Triton X-100 in PBSG, along with chicken anti-GFP or guinea pig-anti PV, and incubated on the cells overnight at 4˚C. The next day, the cells were processed as for confocal imaging, but including only the appropriate AlexaFluor488 secondary. 90 nm gold nanoparticles (Cytodiagnostics) were added at 1:10 dilution for 10 minutes before imaging as fiducials for drift and chromatic aberration correction.

| Primary antibodies | Source | RRID | Stock concentration | Dilution |
| --- | --- | --- | --- | --- |
| Polyclonal chicken anti-GFP | Thermo Fisher A10262 | AB_2534023 | 1 mg/mL | 1:500 |
| Guinea pig anti-parvalbumin antiserum | Swant GP72 | AB_2665495 | 1 mg/mL | 1:500 |
| Monoclonal mouse IgG2A anti-PSD95 (clone K28/43) | Neuromab 75-028 | AB_2877189 | 0.5 mg/mL | 1:80 |
| Polyclonal rabbit anti-Munc13 | Synaptic Systems 126103 | AB_887733 | 0.5 mg/mL | 1:250 |

| Secondary reagents | Source | RRID | Stock concentration | Dilution |
| --- | --- | --- | --- | --- |
| Donkey anti-chicken AlexaFluor488 | Jackson ImmunoResearch 703-545-155 | AB_2340375 | 1.25 mg/mL | 1:500 |
| Donkey anti-guinea pig AlexaFluor488 | Jackson ImmunoResearch 706-545-148 | AB_2340472 | 1.25 mg/mL | 1:500 |
| Goat anti-mouse IgG2a | Jackson ImmunoResearch 115-005-206 (conjugated to Cy3B in-house) | AB_2338462 | ~1.25 mg/mL | 1:500 |
| Donkey anti-rabbit AlexaFluor647 | Jackson ImmunoResearch 711-605-152 | AB_2492288 | 1.25 mg/mL | 1:500 |
| FluoTag-XM-QC anti-mouse IgG kappa light chain sdAb (clone 1A23) + docking site F1 | Massive Photonics (custom) | | 5 μM | n/a |
| FluoTag-XM-QC anti-rabbit IgG sdAb (clone 10E10) + docking site F3 | Massive Photonics (custom) | | 5 μM | n/a |
| ChromPure Rabbit IgG, Fc fragment | Jackson ImmunoResearch 011-000-008 |  | n/a | n/a |

### Confocal microscopy

Confocal images were acquired on a Nikon TI2 Eclipse inverted microscope equipped with a piezo unit for Z control (ASI), a Nikon Apo TIRF 60x/1.49 NA objective and a Dragonfly confocal unit (Andor). Excitation laser light (488, 561, or 638 nm) from an Andor ILE, flattened by an Andor Beam Conditioning Unit, was passed to the sample by a 405/488/561/640 quadband polychroic (Chroma). Emission light was passed through an appropriate bandpass filter (ET525/50, ET600/50 (Chroma), or Em01-R442/647 (Semrock), for AlexaFluor488, Cy3B, and AlexaFluor647 emission, respectively) and collected on a Zyla 4.2 sCMOS camera (Andor). Cells of interest were imaged with 10 µm thick confocal z-stacks with 0.3 µm z-steps at 50% laser power (∼1-2 W/cm^2^) with 200 ms exposure, with each channel imaged sequentially. Reference bead stacks for chromatic aberration correction were acquired by immobilizing 100 nm TetraSpeck beads (Invitrogen) on a poly-L-lysine-coated coverslip and imaging z-stacks as above.

### Confocal analysis

Most confocal image processing was performed with custom macros in FIJI (60), and the processing steps are summarized in Figure S1. First, a reference translation mask for chromatic aberration correction was generated from the TetraSpeck bead stacks using the NanoJ Core (61) plugin *Register Channels – Estimate*, then experimental z-stacks were converted to maximum intensity projections and chromatic aberrations corrected using NanoJ Core *Register Channels – Apply.* Next, each image was cropped to clean areas of GFP or PV signal, background subtracted by the lowest 1% pixel value per channel, and the GFP or PV image manually thresholded to select dendrites. The dendrite masks were cleaned up in two steps. First, the binary was smoothed with *Process>Binary>Dilate* 2x followed by *Close Holes*, then, small background particles in the mask were identified by *Analyze>Analyze Particles* with a size limit of 0-10 pixels, and each identified spot was removed from the mask by an XOR operation against the entire selection. The binary image was cropped to the resulting ROI and used as the dendrite mask. Putative synaptic puncta were identified in each channel with permissive settings of the plugin SynQuant (31; z-score 10, minfill 0.65, whratio 6, min size 10, max size 200), which in our dataset identified many real puncta but also some generally large background spots. These spots were refined through a series of filtering steps 1) postsynaptic ROIs were removed if they did not overlap with the dendrite mask; 2) as we found that SynQuant worked well to identify puncta but less well to identify puncta size in our dataset, the remaining postsynaptic ROIs were dilated by 2 pixels to encompass the entire puncta then thresholded to 40% of the maximum intensity within the ROI to better capture the entire puncta; 3) presynaptic ROIs were removed if they did not overlap with the postsynaptic puncta from step 2; 4) remaining presynaptic ROIs were dilated and thresholded as in step 2; 5) postsynaptic ROIs were removed if they did not overlap with the presynaptic ROIs from step 4. PSD-95 and Munc13 puncta mean and integrated intensities as well as area were finally measured from the background-subtracted images in the remaining post-and pre-synaptic ROIs, respectively. Finally, pre and postsynaptic puncta were paired, using a custom script in MATLAB, by, for each PSD-95 ROI centroid, identifying the mutually closest Munc13 ROI centroid within a search radius of 500 nm. Outliers with area greater than 0.5 µm^2^ were removed from the dataset as puncta larger than this were frequently visibly mis-segmented. Data were processed in R and intensities normalized to the mean of the Ex→Ex group, per week, to control for variance in staining and imaging across culture weeks.

### Single-molecule microscopy

3D DNA-PAINT images were acquired on a custom microscope built around an RM21 base (Mad City Labs), equipped with an infinity-corrected Olympus UPlanApo60x OHR/1.5 NA objective and stage piezo nanopositioner, as well as both micromirror and dichroic-based TIRF pathways. Excitation lasers (405, 488, 561, 638, and 785 nm; Oxxius) are separately expanded 10x by an *f*=25 mm asphere and *f*=250 mm achromatic doublet lens pair, cleaned up by a narrow bandpass filter, and cropped to 8 mm by an adjustable iris, then are combined into the same path by a series of longpass dichroics. The 785 nm laser additionally passed through an optical isolator before beam expansion, and a 25 µm pinhole just after the asphere to produce a higher quality beam for focus lock. To achieve focus lock, the 785 nm laser is passed through a T660lpxr dichroic and directed to the micromirror TIRF pathway of the RM21, which directs the beam to TIR and catches the exit beam on the far side of the objective. The exit beam is passed through a cleanup filter (ET780lp) and passed to a QPD such that deflections in the beam position due to focus shifts are compensated in a closed-loop system with the stage nanopositioner to maintain sample focus with high fidelity and responsiveness. The 405, 488, 561 and 638 nm lasers are reflected by the T660lpxr dichroic and directed to the dichroic TIRF module, which directs and focuses the beams off-center in the back aperture of the objective via a ZT405/488/561/640 rpvc2 quadband polychroic to achieve an adjustable illumination angle of the sample from epi to TIR. The micromirror and dichroic TIRF modules direct the laser to orthogonal positions on the objective back aperture for simultaneous imaging and focus lock. Emission light is focused by an *f*=300 mm achromatic doublet onto an adjustable rectangular iris, and is split in the MadView (MadCity Labs), in which a series of longpass dichroics reflect emission from 488, 561 and 640 excitation through separate adjustable *f*=225 mm lenses, allowing separate control of chromatic aberrations for each imaging channel. The emission lines are recombined by paired dichroics and focused onto a Prime 95b sCMOS camera (Photometrics) with an *f*=250 mm lens, first passing through an *f*=500 mm cylindrical lens to provide astigmatism for 3D imaging. The recombination of emission lines was done such that two-color simultaneous imaging was achieved by separating the parts of the camera chip receiving each color. The system achieves a final magnification of 110x, yielding a pixel size of 100 nm.

### Single-molecule imaging

GFP or PV-positive cells were identified with low power 488 nm laser. F1-Atto643 (for PSD-95) and F3-Cy3B (for Munc13) imagers were diluted in imaging buffer (1x PBS pH7.4 + 500 mM NaCl + PCA/PCD/Trolox oxygen scavengers; 59) to 0.25 nM each and added to the sample, which was allowed to equilibrate for at least 10 minutes to reduce drift. Then, 40,000 frames were acquired with 150 ms exposure, with laser power densities at the sample of 0.10 kW/cm^2^ for the 638 nm laser and 0.059 kW/cm^2^ for the 561 nm laser. The 785 nm laser was set to 50% power and used for focus lock. Imaging buffer was refreshed between regions.

### Single-molecule localization

Molecule locations were determined using the Super-resolution Microscopy Analysis Platform, SMAP (62). Z stacks of the 3D PSF of 100 nm TetraSpeck beads on coverglass were captured with 10 nm step size and a standard curve relating PSF characteristics to z position, from which z position of molecules was derived, was generated in SMAP using the *calibrate3DsplinePSF* plugin. Parameters were kept as default with the following exceptions: 1) the 3D modality was set as *global 2 channel* and *focus as z ref* was toggled to treat the in-focus PSF as the 0 nm position in z; 2) *Global Fit Parameters* were modified to set *main Ch* as *u/l* and *right-left* with *Split (pix)* set to the distance in x in pixel units that bisected the image – this, in accordance with the simultaneous two-color imaging described above, ensures that the same beads formed the basis of each color’s individual calibrations. Beads were selected on one side of the image using *Spatially resolved calibration: Interactive ROI, rectangular, right-left* with *no mirror*. To enable chromatic aberration correction and overlay of channels (see below), 5-10 regions of in-focus TetraSpeck beads were imaged without astigmatism for 100 frames and localized using SMAP. Images were loaded using *TifLoader*. *PeakFinder* settings were left at default and *PSF free* fitter was used with ROI size 7. For 3D experimental images, *PeakFinder* and *Fitter* settings were adjusted such that *DoG = 2.5*, *cutoff = 2*, *ROI size = 15*, and the *spline* fitter was used, adjusting the RI mismatch to 0.83 as appropriate for the system and loading the previously calculated 3D calibration.

### Single-molecule analysis

All analysis was conducted using custom routines in MATLAB (Mathworks) which relied, in part, on command line calls to Picasso (63) for specific functions.

#### Processing of super-resolution images

To correct chromatic aberrations between the channels, the localizations from the 2D bead images described above were first bisected and overlaid, followed by a nearest neighbor approach to match localized beads between the channels. A transformation matrix was subsequently generated using the *fitgeotrans* function in MATLAB with the input (‘polynomial’,2). This transformation was applied to the experimental data following cross-correlation-dependent drift correction, which was conducted using Picasso Render’s *Undrift by RCC* function. The two channels were then more precisely aligned using a cross-correlation to align fiducial markers and then treated individually hereafter until noted otherwise. Poorly fit molecules were eliminated if their: localization precision was greater than 20 nm, PSF standard deviation was greater than 2 pixels, photon count was smaller than the mode of the whole-field histogram, or relative log likelihood was less than the first shoulder in the histogram (∼-1.5). This resulted in a median xy localization precision for Munc13-1 of 2.35 nm in the Ex→Ex synapses and 2.54 nm in the Ex→PV synapses and a median xy localization precision for PSD-95 of 6.52 nm in the Ex→Ex synapses and 6.94 nm in the Ex→PV synapses. Localizations that persisted over multiple frames were merged using Picasso Render’s *Link localizations* function, with max distance of 30 nm and max transient dark frames of 5. We noticed that the SMAP localization resulted in some molecules localized at an artificial extreme ceiling or floor in z, whose specific value depended on the calibration calculated in SMAP; these were eliminated from the analysis on a per image basis.

#### Identifying putative synapses

To automatically segment objects from the image, the Picasso Render *Clustering>DBSCAN* function was used with radius of 30 nm and minimum density of 5. Identified objects were filtered on the mean and standard deviation of the frames in which localizations within each object were present – this helped avoid inclusion of leftover gold fiducials (high standard deviation of frame number) or any clusters that were formed by nonspecific binding and localization of individual imager strands (low/high mean and low standard deviation of frame number). Clusters were further filtered by extracting PSD-95 clusters whose 2D projection overlapped with at least 1 localization of Munc13-1, using the *alphaShape* and *inShape* functions in MATLAB. The resultant clusters were then used to similarly filter the Munc13-1 clusters. Both protein species were then treated as if from the same population of molecules and were segmented into putative synapses using MATLAB’s *dbscan* function with epsilon of 30 nm and minpts of 4. As a quality control measure, manual inspection of the resulting putative synapses enabled manual segmentation where DBSCAN had failed to separate two synaptic clusters. To filter down to the best sampled and well-segmented synapses, PSDs that contained fewer than 20 localizations, or whose long/short axis ratio was greater than 4, or whose area was less than 1.5 pixels or larger than 20 pixels were removed. The remaining putative synapses were then judged for quality based on sampling density, corresponding presence and shape between both Munc13-1 and PSD-95 clusters, and z spread of localizations. Synapses were sorted for cis-and/or trans-synaptic analyses based on these qualities and putative outlier localizations (which could impact synaptic volume and shape-based analyses) were removed by calculating the mean x, y, and z position of each point cloud and keeping only those localizations that fell within 2 standard deviations.

#### Quantitative analysis of synaptic nanostructure

Auto-and cross-correlation analyses were conducted as described previously (10). Nanocluster detection relied on the *dbscan* function in MATLAB. The pairwise distance between each localization was computed and input into the *dbscan* function, along with an epsilon and minpts value that is unique to each point cloud, and the optional inputs (‘Distance’,’precomputed’). Epsilon for each point cloud was determined first by calculating the mean minimal distance for each point within the point cloud and multiplying by an empirically determined factor that was specific to each protein species (30 random point clouds tested to determine most representative factor yielded 1.6 for Munc13-1 and 2.7 for PSD-95). Local density heat maps shown in Figures 2-4 were generated using the calculated epsilon for each synapse and thus represent the input to DBSCAN. The mean number of neighboring points within epsilon for each localization was calculated and minpts was set as 1.96 standard deviations above the result, rounded down. Synaptic and nanocluster volumes were calculated using either the entire synaptic point cloud or just the localizations assigned to a specific NC, respectively, by using the *volume* and *alphaShape* functions with an alpha radius of 150 nm. The percentage of synaptic volume occupied by NCs was calculated by summing the individual volumes of each NC in each synapse and dividing by the overall volume of the matched synaptic compartment. Auto-and cross-enrichment analyses were calculated as described previously (10, 14), with one exception. The normalization in both analyses was conducted to a pseudorandom distribution of the same number of measured localizations whose only constraint was to guarantee one localization per bin – this was done to eliminate *NaN* and *Inf* outputs that would mask data in the real distribution due to 0s appearing in the randomized one. This strategy could result in individual bins that were normalized to a value of 1 having an outsized impact on the shape of the curve. To mitigate this possibility, the auto-and cross-enrichment curves were smoothed by first detecting outliers in each distance bin using a ROUT of 0.1% in PRISM (GraphPad), then calculating the maximum non-outlier value in each bin, and finally replacing the outlier values in each bin with the corresponding value. This maintains the influence of high-density regions on the curve but tempers inflation due to random chance. Prior to the cross-enrichment, the PSD-95 point cloud was shifted onto the Munc13-1 point cloud based on the cross-correlation of their shape, agnostic to any subsynaptic heterogeneity (10, 64).The percent of Munc13-1 NCs aligned with PSD-95 was calculated using the same base calculation of the cross enrichment, which yields a relative enrichment of the opposing protein within binned distances from the peak of a given NC. These values were averaged within a specific window. In parallel, a similar calculation was conducted on 50 random distributions of the opposing protein, bounded by the original cluster’s shape and matching the original number of localizations. A NC was considered aligned if the average value was greater than 1.96 standard deviations away from the mean of the random distributions. The range of the specific window depended on the analysis. In Fig 4G, it was within 60 nm of the peak. In Fig 4H, an iterative range was used in increments of 10 nm from 20-150 nm. In Fig 4I, a window of set width (30 nm) was used, but the binned distances used in the calculation were shifted within a range of 10-150 nm from the NC peak.

#### Statistical analysis

Statistical analysis was conducted using GraphPad PRISM. Data were tested for normality using D’Agostino & Pearson, Shapiro-Wilk, and Kolmogorov-Smirnov tests. Data that passed normality checks were tested using two-tailed t-tests if variance was equal between groups or Welch’s t-test if not. Data that did not pass these checks were tested with two-tailed Mann-Whitney tests. No statistical methods were used to predetermine sample size. Experimenters were blinded during super-resolution analyses.

## Acknowledgements

This work was supported by F31 MH117920 and T32 GM008181 to P.A.D., F32 MH119687 to A.D.L, and R37 MH080046 and R01 MH119826 to T.A.B. We thank the members of the Blanpied Laboratory for rigorous discussion and critical evaluation of the manuscript, and Minerva Contreras for technical assistance.

---

**Figure S1.** Visual outline of image processing and quantification steps for confocal measurements in Figure 1.

Details are in Materials and Methods. **1.** Images are pre-processed for analysis. Confocal z-stacks are maximum intensity projected, corrected for chromatic aberrations between channels, cropped to an area of interest, and background subtracted. **2.** The dendrite mask is created to isolate synapses to the cell of interest. The dendrite signal is thresholded and binarized, and the binary adjusted. Small spots arising from off-cell background are selected then removed by an XOR operation against the entire selection, resulting in a clean binary. **3 and schematic.** Synaptic puncta are identified on the cell of interest. Schematic shows which puncta are removed in each filtering step shown in 3; magenta are postsynaptic puncta and yellow presynaptic. ROIs are identified by SynQuant in both channels. Then, postsynaptic ROIs are filtered to those that overlap with the dendrite mask and adjusted to the size of the actual puncta. Presynaptic ROIs are filtered to those that overlap with the postsynaptic ROIs, adjusted for size, and finally postsynaptic ROIs are filtered to those that overlap with the presynaptic ROIs, leaving a set of putative “synaptic” ROIs. ROIs are confirmed as synaptic if they have a mutually closest neighbor within a 500 nm maximum distance, then are quantified.

