## Supplemental Figure 1 for "Differential nanoscale organization of excitatory synapses onto excitatory vs inhibitory neurons"

### Figure S1

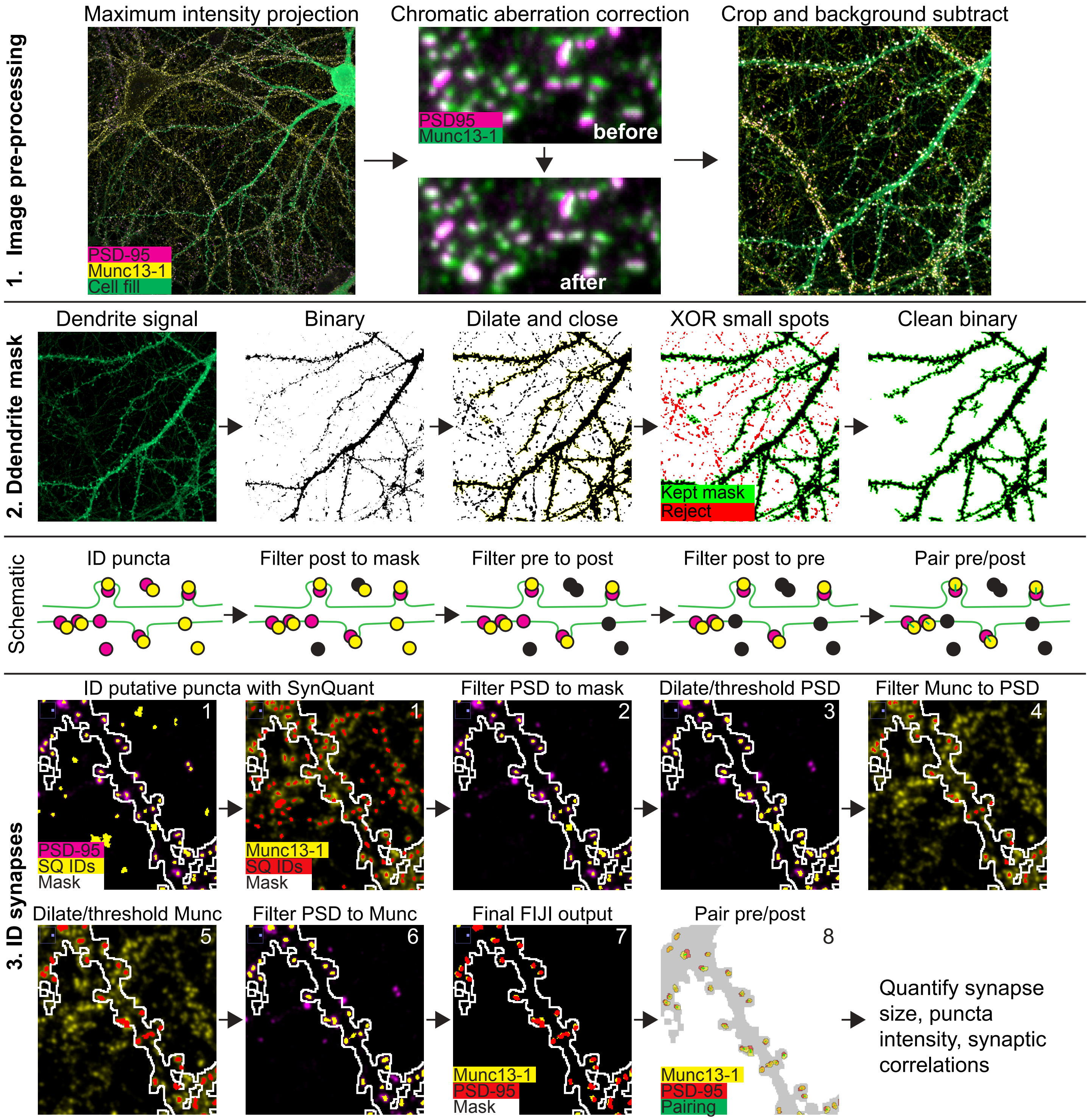
1. Image pre-processing
Maximum intensity projection
PSD-95
Munc13-1
Cell fill
Chromatic aberration correction
PSD95
Munc13-1
before
after
Crop and background subtract
2. Ddendrite mask
Dendrite signal
Binary
Dilate and close
XOR small spots
Kept mask
Reject
Clean binary
Schematic
ID puncta
Filter post to mask
Filter pre to post
Filter post to pre
Pair pre/post
3. ID synapses
ID putative puncta with SynQuant
1
PSD-95
SQ IDs
Mask
1
Munc13-1
SQ IDs
Mask
Filter PSD to mask
2
Dilate/threshold PSD
3
Filter Munc to PSD
4
Dilate/threshold Munc
5
Filter PSD to Munc
6
Final FIJI output
7
Munc13-1
PSD-95
Mask
Pair pre/post
8
Munc13-1
PSD-95
Pairing
Quantify synapse size, puncta intensity, synaptic correlations

**Figure S1.** Visual outline of image processing and quantification steps for confocal measurements in Figure 1.

Details are in Materials and Methods. **1.** Images are pre-processed for analysis. Confocal z-stacks are maximum intensity projected, corrected for chromatic aberrations between channels, cropped to an area of interest, and background subtracted. **2.** The dendrite mask is created to isolate synapses to the cell of interest. The dendrite signal is thresholded and binarized, and the binary adjusted. Small spots arising from off-cell background are selected then removed by an XOR operation against the entire selection, resulting in a clean binary. **3 and schematic.** Synaptic puncta are identified on the cell of interest. Schematic shows which puncta are removed in each filtering step shown in 3; magenta are postsynaptic puncta and yellow presynaptic. ROIs are identified by SynQuant in both channels. Then, postsynaptic ROIs are filtered to those that overlap with the dendrite mask and adjusted to the size of the actual puncta. Presynaptic ROIs are filtered to those that overlap with the postsynaptic ROIs, adjusted for size, and finally postsynaptic ROIs are filtered to those that overlap with the presynaptic ROIs, leaving a set of putative “synaptic” ROIs. ROIs are confirmed as synaptic if they have a mutually closest neighbor within a 500 nm maximum distance, then are quantified.
